# Individual-Specific fMRI-Subspaces Improve Functional Connectivity Prediction of Behavior

**DOI:** 10.1101/515742

**Authors:** Rajan Kashyap, Ru Kong, Sagarika Bhattacharjee, Jingwei Li, Juan Zhou, Thomas Yeo

## Abstract

There is significant interest in using resting-state functional connectivity (RSFC) to predict human behavior. Good behavioral prediction should in theory require RSFC to be sufficiently distinct across participants; if RSFC were the same across participants, then behavioral prediction would obviously be poor. Therefore, we hypothesize that removing common resting-state functional magnetic resonance imaging (rs-fMRI) signals that are shared across participants would improve behavioral prediction. Here, we considered 803 participants from the human connectome project (HCP) with four rs-fMRI runs. We applied the common and orthogonal basis extraction (COBE) technique to decompose each HCP run into two subspaces: a common (group-level) subspace shared across all participants and a subject-specific subspace. We found that the first common COBE component of the first HCP run was localized to the visual cortex and was unique to the run. On the other hand, the second common COBE component of the first HCP run and the first common COBE component of the remaining HCP runs were highly similar and localized to regions within the default network, including the posterior cingulate cortex and precuneus. Overall, this suggests the presence of run-specific (state-specific) effects that were shared across participants. By removing the first and second common COBE components from the first HCP run, and the first common COBE component from the remaining HCP runs, the resulting RSFC improves behavioral prediction by an average of 11.7% across 58 behavioral measures spanning cognition, emotion and personality.

**Highlights:**

- We decomposed rs-fMRI signals into common subspace & individual-specific subspace
- Common subspace is shared across all Human Connectome Project (HCP) participants
- Common subspaces are different across runs, suggesting state-specific effects
- Individual-specific subspaces are unique to individuals
- Removal of common subspace signals improve behavioral prediction by 11.7%

## Introduction

Mapping from brain to behavior in individuals is a crucial step in developing imaging-based biomarkers with real-world utilities. With the widespread availability of large-scale resting-state functional magnetic resonance imaging (rs-fMRI) datasets, there has been significant interest in using resting-state functional connectivity as a predictive fingerprint of human behavior (Finn et al., 2015; Rosenberg et al., 2016). Resting-state functional connectivity reflects synchrony between brain regions present during rest and has been widely utilized to provide insights into the intrinsic architecture of the human brain (Biswal et al., 1995; Fox and Raichle, 2007; Buckner et al., 2013). The brain functional architecture measured during the resting-state is similar (although not the same) during task states, suggesting the relevance of resting-state functional connectivity to brain function and cognition (Smith et al., 2009; Mennes et al., 2010; Yeo et al., 2015a; Tavor et al., 2016). Consequently, resting-state functional connectivity has been widely utilized to predict behavioral measures, ranging from cognition to personality (Hampson et al., 2006; Smith et al., 2015; Dubois et al., 2018; Bertolero et al., 2018).

Successful behavioral prediction requires functional connectivity to be distinct across individuals, while retaining key features of an individual (Finn and Constable, 2016). For example, if the functional connectivity patterns of all participants were the same, then behavioral prediction could not possibly work. Therefore, fMRI signals that are present (or shared) across participants should theoretically not be useful for prediction. In fact, the shared signals might confuse the prediction algorithm, leading to worse prediction performance. Here, we investigated whether removing fMRI signals that are common across individuals might improve behavioral prediction.

More specifically, we applied the common orthogonal basis extraction (COBE) algorithm (Zhou et al., 2016a; Zhou et al., 2016b) to rs-fMRI data from the Human Connectome Project (HCP; Van Essen et al., 2012; Smith et al., 2013). The COBE algorithm was originally developed to project “multi-block” data (collection of matrices) into a common subspace shared by all blocks and block-specific subspaces. The number of components spanning each subspace is specified by the user. The rs-fMRI data of an individual could be thought of a block, so application of COBE to the rs-fMRI of all participants decomposed each participant’s fMRI signals into a linear sum of a number of common COBE components (shared by all participants) and a number of individual-specific COBE components. Our hypothesis is that removing the common (group-level) COBE components from the rs-fMRI data might yield an improved predictive fingerprint of human behavior.

Conceptually, it is worth distinguishing our work from the vast literature investigating trait-level and state-level aspects of functional connectivity (Shirer et al., 2012; Cole et al., 2014; Krienen et al., 2014; Meija et al, 2015; Yeo et al., 2015b; Wang et al., 2016; Gratton et al., 2018; Green et al., 2018; Kong et al., 2018). Here, our goal was to remove rs-fMRI signals common across participants, which might include common state-level effects (e.g., arising from participants undergoing the same experimental protocol), but also trait-level effects shared across all participants (e.g., all HCP participants are young adults).

Cognizant of the fact that there might be inter-run variation (state-level effects) across the four runs of the HCP data (Bijsterbosch et al., 2017), the COBE algorithm was applied to each HCP run independently to explore if the common COBE components were similar across runs. We then evaluated whether functional connectivity computed using the individual-specific fMRI signals can improve prediction of 58 behavioral measures across cognition, personality and emotion.

## Methods

### Overview

COBE was applied to preprocessed rs-fMRI data of 803 subjects from the Human Connectome project (HCP). Three variants of COBE emerged and the individual-subspace functional connectivity derived from these variants were considered for prediction of 58 behavioral measures. Prediction accuracies with and without COBE were assessed.

### Rs-fMRI data

The HCP S1200 release comprises a multi-modal collection of data across behavioral, structural MRI, rs-fMRI, and MEG paradigms from healthy adults (Van Essen et al., 2012; Smith et al. 2013). All imaging data were collected on a custom-made Siemens 3T Skyra scanner using a multiband sequence. The MRI and behavioral data were collected on two consecutive days. During the resting-state scan, participants are instructed to fixate their eyes on a projected bright cross-hair on a dark background. There were two rs-fMRI sessions. Each rs-fMRI session consisted of two runs. For convenience, we will refer to the two rs-fMRI runs (obtained on the first day) as run 1 and run 2. We will refer to the two rs-fMRI runs (obtained on the second day) as run 3 and run 4. Each rs-fMRI run was acquired in 2mm isotropic resolution with a TR of 0.72 seconds for a total of 1200 frames lasting 14 minutes and 33 seconds (Van Essen et al., 2012; Smith et al., 2013).

### Preprocessing

We utilized the MSMAll ICA-FIX data on fs_LR32K surface space (HCP S1200 manual; Glasser et al. 2013; Griffanti et al., 2014; Salimi-Khorshidi et al. 2014) from 1094 participants. However, some studies have pointed out that ICA-FIX does not completely eliminate global head-motion artefacts and recommended further nuisance regression (Burgess et al., 2016; Siegel et al., 2016; Kong et al., 2018; Li et al., 2019). More specifically, framewise displacement (FD; Jenkinson et al., 2002) and root-mean-square of voxel-wise differentiated signal (DVARS) (Power et al., 2012) were estimated using fsl_motion_outliers. Volumes with FD > 0.2mm and DVARS > 75, as well as uncensored segments of data lasting fewer than 5 contiguous volumes were flagged as outliers. Nuisance regression with regressors consisting of a global signal, six motion parameters, averaged ventricular signal, averaged white matter signal, and their temporal derivatives (18 regressors in total) were performed. When performing nuisance regression, outlier volumes were ignored in the computation of the regression coefficients. A bandpass filter (0.009 Hz ≤ f ≤ 0.08 Hz) was then applied to the data. BOLD runs with more than half the volumes flagged as outliers were completely removed. Consequently, 82 subjects had all runs removed and were thus not considered further.

Preprocessed rs-fMRI time courses were averaged within each of 400 cortical parcels (Schaefer et al., 2017) and 19 subcortical regions (brain stem, accumbens, amygdala, caudate, cerebellum, diencephalon, hippocampus, pallidum, putamen, and thalamus; Fischl et al., 2002). Therefore, there were 419 regions in total, resulting in a 419 x 1200 matrix of rs-fMRI time courses for each run of each subject.

### Behavioral data

We considered a set of 58 behavioral measures across cognition, personality and emotion (Table S1; Kong et al., 2018). We restricted our analyses to participants, who had all four runs survived the quality control procedure and all 58 behavioral measures, resulting a final set of 803 participants.

### Common Orthogonal Basis Extraction (COBE)

For more details about the COBE algorithm, we refer readers to previously published papers (Zhou et al., 2016a; Zhou et al., 2016b). Here we briefly describe how COBE was applied to rs-fMRI data in this study.

Given that within-subject differences have been reported across the four HCP runs (Bijsterbosch et al., 2017), COBE was applied to the four runs separately. In other words, COBE was applied to the first runs of all subjects, the second runs of all subjects, the third runs of all subjects and finally, the fourth runs of all subjects.

For ease of explanation, let us consider the first run of all subjects. As explained previously, the first run of a subject is represented as a 419 × 1200 matrix. Let *S*_*n*_ denote the 419 × 1200 rs-fMRI matrix of the *n*-th subject. As illustrated in Figure 1A, COBE seeks to decompose *S*_*n*_ into

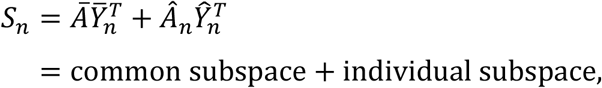

where *Ā* is a 419 × C matrix representing the common subspace shared across all subjects. C is the number of components spanning the common subspace, and is defined a priori by the user. Thus, COBE assumes spatial correspondence, but not temporal correspondence across subjects. Indeed, each column of *Ā* can be visualized as a spatial map (see Figure 2 in Results). 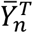 is a C × 1200 matrix of subject-specific time courses (of the *n*-th subject) associated with the common space. 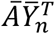 is a 419 × 1200 matrix representing the projection of the *n*-th subject’s rs-fMRI time courses onto the common subspace, while 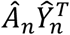 is a 419 × 1200 matrix representing the projection of the *n*-th subject’s rs-fMRI time courses onto the individual-specific subspace. Figure 1B illustrates *S*_*n*_ (red), 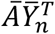(black) and 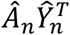 (green) for a random HCP subject with C = 1.

**Figure 1.**
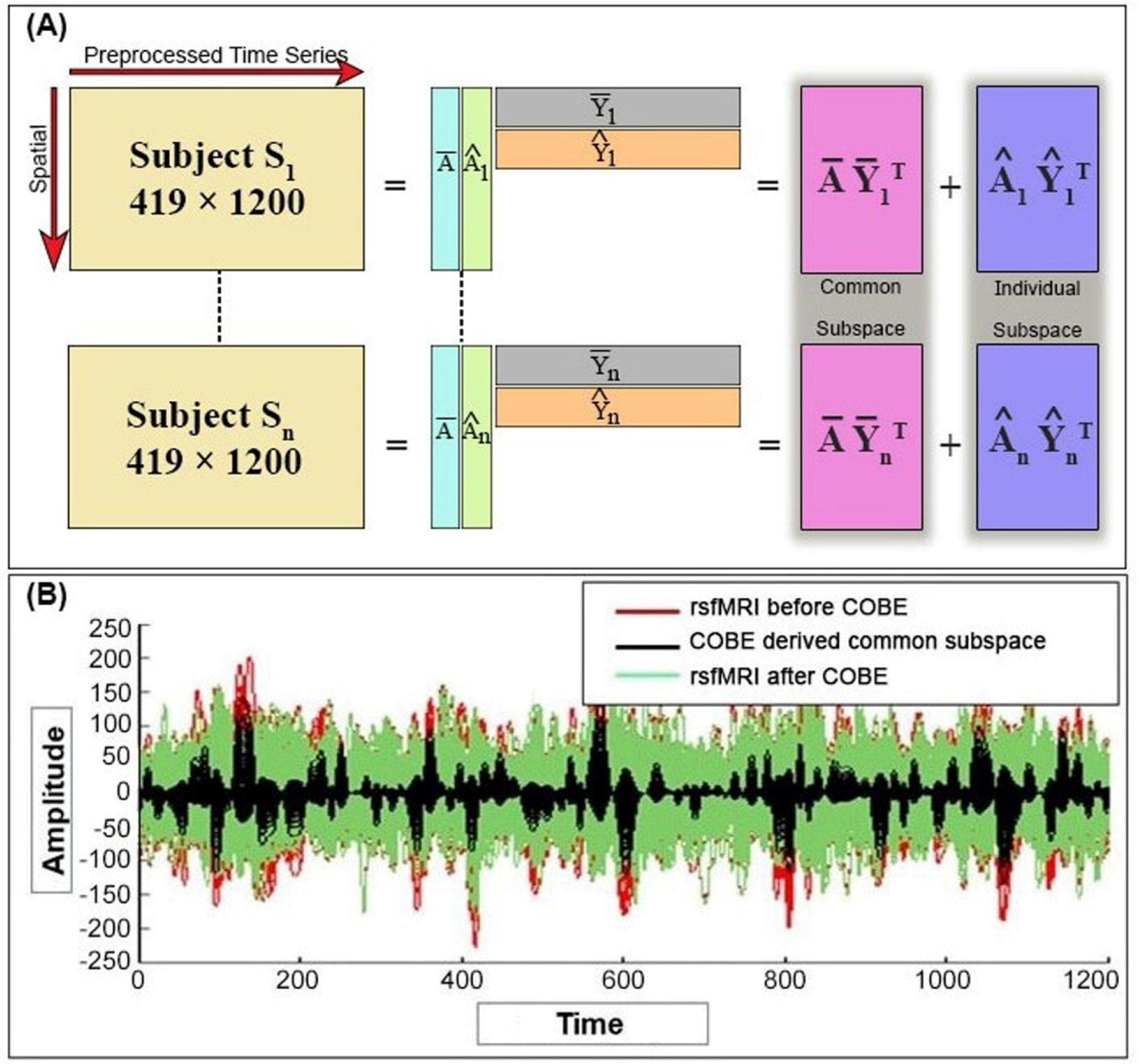
Illustration of Common Orthogonal Basis Extraction (COBE). (A) COBE applied to one rs-fMRI run of 803 HCP participants. COBE projects the rs-fMRI data (*S*_*n*_) of an HCP individual onto a common subspace 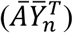 and individual-specific subspace 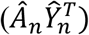. The common subspace (*Ā*) is shared across all subjects. The number of components C spanning the common subspace (number of columns of *Ā*) is a user-specified parameter. (B) The original signals (red), common-subspace signals (black; 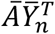) and individual-subspace signals (green; 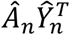) are shown for a random HCP subject

**Figure 2.**
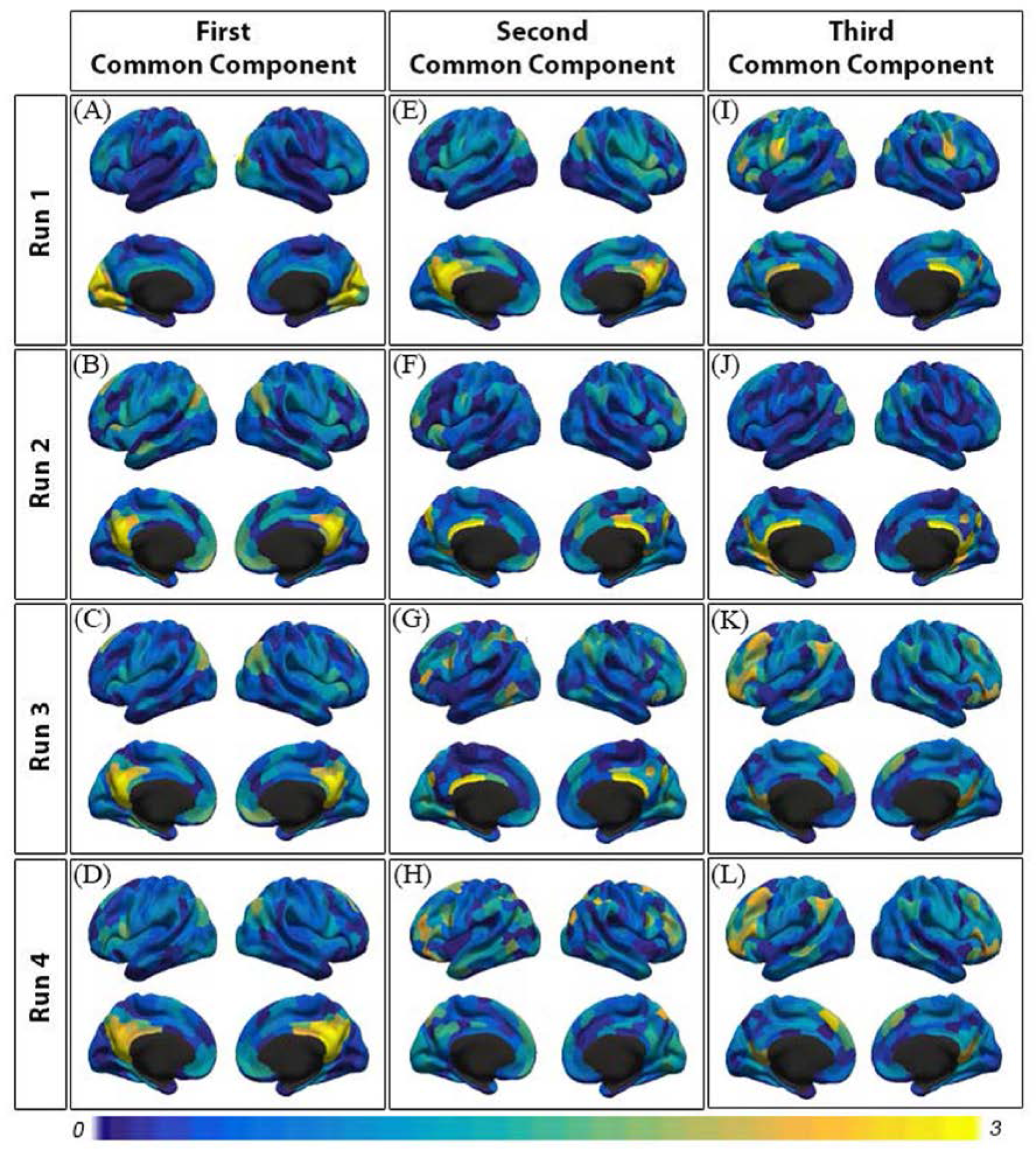
Spatial map of three common components shared across subjects (N = 803) in each of the four rs-fMRI runs. Observe that the first component of the first run (Figure 2A) was unique to only that run. Instead, the second component of the first run and the first component of the remaining runs were highly similar (r = 0.83). The third component of the first run and the second component of the remaining runs were also similar (r = 0.42). We note that very similar components were obtained if the rs-fMRI time courses were variance normalized before COBE was applied.

### Three variants of COBE

An important parameter is the number of common components C. A useful property of COBE is that if COBE was applied twice (sequentially) with C = 1, the two common components will be (in practice) the same as the common components obtained by applying COBE once with C = 2. Here, we applied COBE extracting to each of the four HCP runs, sequentially extracting components up to three components per run, yielding 4 × 3 = 12 common components.

As will be elaborated in Figure 2 of the Results section, we found that the first common component of the first run was unique to the run, while the second common component of the first run was highly similar to the first common components of the remaining runs. On the other hand, the third common component of the first run was also similar to the second common components of the remaining runs. These results motivated three variants of COBE for subsequent analyses: (i) COBE-1000, where COBE was only applied to the first run (C = 1) and the common subspace spanned by the common component was removed from the first run, (ii) COBE-2111, where COBE was applied to the first run (C = 2) and the remaining runs (C = 1), and the common subspace spanned by the common components were removed from the respective runs, and (iii) COBE-3222, where COBE was applied to the first run (C = 3) and the remaining runs (C = 2), and the common subspace spanned the common components were removed from the respective runs. Finally, along with three variants of COBE, we also considered “NO-COBE”, where COBE was not utilized at all.

### Behavioral prediction with and without COBE

For each variant of COBE, 419 × 419 RSFC (Pearson’s correlation) matrix was computed based on the individual-subspace signals (419 × 1200) of each run of each subject. The correlation matrices were then averaged across all the four runs of each subject. For example, in the case of the variant COBE-2111, the final correlation matrix of a subject was computed in the following way: (i) for the first run, the correlation matrix was computed after two common-subspace components were removed, (ii) for the remaining runs, the correlation matrices were computed after one common-subspace component was removed from the corresponding runs, and (iii) the four correlation matrices were averaged to obtain the final functional connectivity matrix of a subject. A 419 × 419 Pearson’s correlation matrix was also computed for the “NO-COBE” condition.

As previously discussed, we considered 58 behavioral measures (Table S1, Supplement). For each behavioral measure, the elastic net (Friedman et al., 2010; Zou and Hastie, 2005) was used to predict subjects’ behavior in a 20-fold nested cross-validation scheme using the functional connectivity matrices obtained from COBE-1000, COBE-2111, COBE-3222 or NO-COBE. For every test fold and each behavioral measure, the remaining 19 folds were used for training and validation. More specifically, certain behavioral measures are known to correlate with motion (Siegel et al., 2016). Therefore, age, sex, and framewise displacement (FD) were regressed from the behavioral measure before elastic net regression. The nuisance regression was performed on the training and validation folds, and the estimated coefficients were then applied to the test fold.

After nuisance regression, the 19 training and validation folds were used for feature selection by selecting the top 50% of functional connections most strongly correlated (positive or negative) with the particular behavioral measure (see HCP MegaTrawl; https://db.humanconnectome.org/megatrawl/HCP820_MegaTrawl_April2016.pdf). The selected features (functional connectivity strength) were then entered into the elastic net regression estimation procedure. There were two hyperparameters associated with the elastic net, which were determined via inner-loop cross-validation of the 19 folds. The optimal hyperparameters were then used for predicting the behavioral measure in the test fold. Accuracy was measured by correlating the predicted and actual behavioral measure across all subjects within the test fold (Finn et al., 2015). Thus, for each behavioral measure, the 20-fold cross-validation yielded 20 prediction accuracies.

When comparing different approaches, the prediction accuracies were averaged across all behavioral measures and then the corrected resampled t-test was utilized (Bouckaert and Frank, 2004; Nadeau and Bengio, 2000). The corrected resampled t-test accounted for the fact that the cross-validation accuracies were not independent across folds.

### Code availability

The code for COBE can be downloaded at http://www.bsp.brain.riken.jp/~zhougx/cifa.html. The code for elastic-net is available freely at https://web.stanford.edu/~hastie/glmnet_matlab/. The Schaefer parcellation is available at https://github.com/ThomasYeoLab/CBIG/tree/master/stable_projects/brain_parcellation/Schaefer2018_LocalGlobal.

## Results

### Overview

COBE was applied to each rs-fMRI run of 803 HCP subjects. The spatial maps of the common (group-level) components from the four runs were then examined. The impact of removing the common components on the resulting RSFC was then investigated. Finally, we explored whether removing the common components from the rs-fMRI data improved behavioral prediction.

### Spatial maps of common COBE components

COBE was applied to the rs-fMRI data of 803 HCP subjects to extract three components (C = 3) that were common across subjects. Common components were extracted from the four rs-fMRI runs independently, yielding a 419 × C matrix Ā for each run. Each column of Ā corresponds to the spatial map of a common COBE component, which is visualized in Figure 2.

The first common COBE component of the first run (Figure 2A) was predominantly focused on the visual cortex, especially the portion of the visual cortex involved in peripheral vison. This component was not found in the remaining runs. Instead, the second common COBE component of the first run and the first common COBE component of the second, third and fourth runs were primarily focused on regions within the default network with particular strong emphasis in the posterior cingulate cortex and precuneus. Indeed, average correlation between the first component of the first run with the first component of the remaining runs was only 0.03. On the other hand, the average correlation between the first component of the first run with the second component of the remaining runs was 0.83. Overall, this suggests the existence of a “common-subspace” component present in the first run of the HCP data, but not present in the remaining runs.

The third common component of the first run and the second common COBE component of the second, third and fourth runs were also similar (r = 0.42), with strong weights on the posterior cingulate cortex and lateral inferior frontal lobe, although the degree of similarity was considerably weaker than the earlier components (discussed in the previous paragraph). For example, the somatomotor face region within the central sulcus exhibited strong spatial weights in the third component of the first run, but not in the second components of the remaining runs. The degree of similarity reduced even more with more components (Figure S1). More specifically, the average correlation between the fourth component of the first run with the third component of the remaining runs was 0.06. Therefore, we did not explore more than three components (e.g., COBE-4333) in subsequent results.

The above observations motivated our considerations of three variants of COBE: COBE-1000, COBE-2111 and COBE-3222 in subsequent analyses. To illustrate the notation, COBE-2111 means that two common components were removed from the first HCP run, while one common component was removed from the remaining three HCP runs.

### Functional connectivity changes arising from removing common COBE components

Figures 3A and 3B show the 19 subcortical and 400 cortical ROIs used to compute the 419 × 419 RSFC matrices. To illustrate the effects of removing common COBE components on the resulting RSFC matrices, Figures 3C and 3D show the RSFC matrices of the first rs-fMRI run (averaged across 803 subjects) before and after removing the first common COBE component. Figure 3E shows RSFC changes from removing the first common COBE component. Since the first common COBE component of the first run was primarily focused on the visual cortex (Figure 2A), removing the first common COBE component largely resulted in RSFC changes associated with the visual network. In particular, there was decreased connectivity within the visual network, decreased connectivity between visual network regions and the somatomotor and dorsal attention networks, as well as increased connectivity between visual network regions and the control and default networks.

Similarly, FC changes from removing the first common COBE component from the second, third and fourth rs-fMRI runs are shown in Figures 4A to 4C. Given that the first component was primarily focused on the posterior cingulate and precuneus (Figures 2B to 2D), the resulting RSFC changes were largely limited to the default network. More specifically, there was decreased connectivity within the default network, as well as increased connectivity between the default and attentional networks (Figures 4A to 4C).

Figure 4D shows FC changes from removing the first two common COBE components from the first rs-fMRI run. Given that the first component was associated with the visual cortex, while the second component was associated with the posterior cingulate and precuneus, the resulting FC changes involved both visual and default networks. Indeed, the resulting FC changes appeared to be a combination of Figure 3E and Figures 4A to 4C.

**Figure 3.**
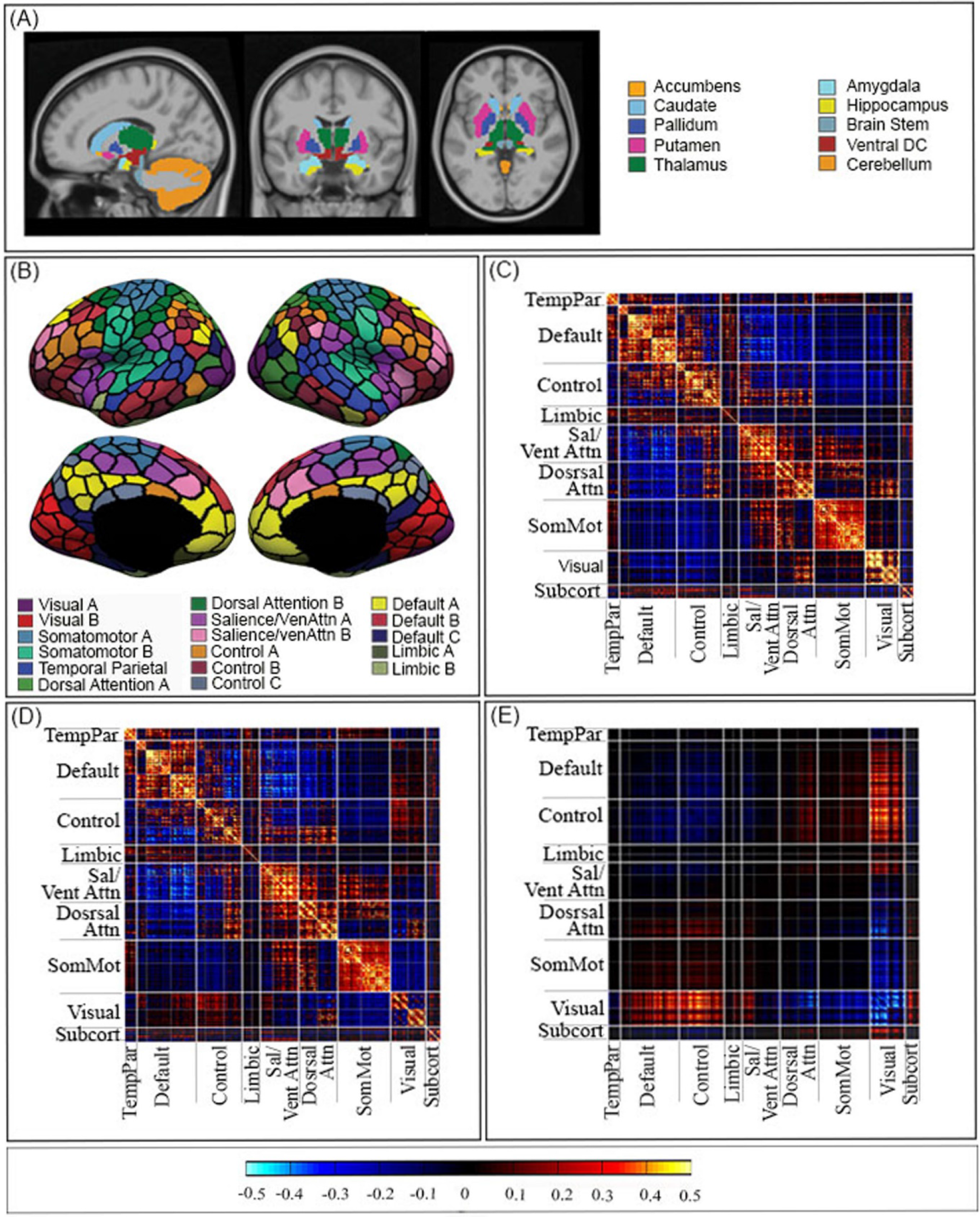
FC changes from removing first COBE component from the first run of the HCP data. 19 subcortical ROIs (Fischl et al., 2002) (B) 400 cortical parcels (Schaefer et al., 2017). Parcel colors correspond to 17 large-scale-networks (Yeo et al., 2011). For visualization, the 17 networks were divided into eight groups (TempPar, Default, Control, Limbic, Salience/Ventral Attention, Dorsal Attention, Somatomotor and Visual). (C) 419 × 419 FC matrix of the first run of the HCP data, averaged across 803 participants. (D) 419 × 419 FC matrix after removing the first common COBE component from the first HCP run. (E) FC difference obtained by subtracting (C) from (D).

**Figure 4.**
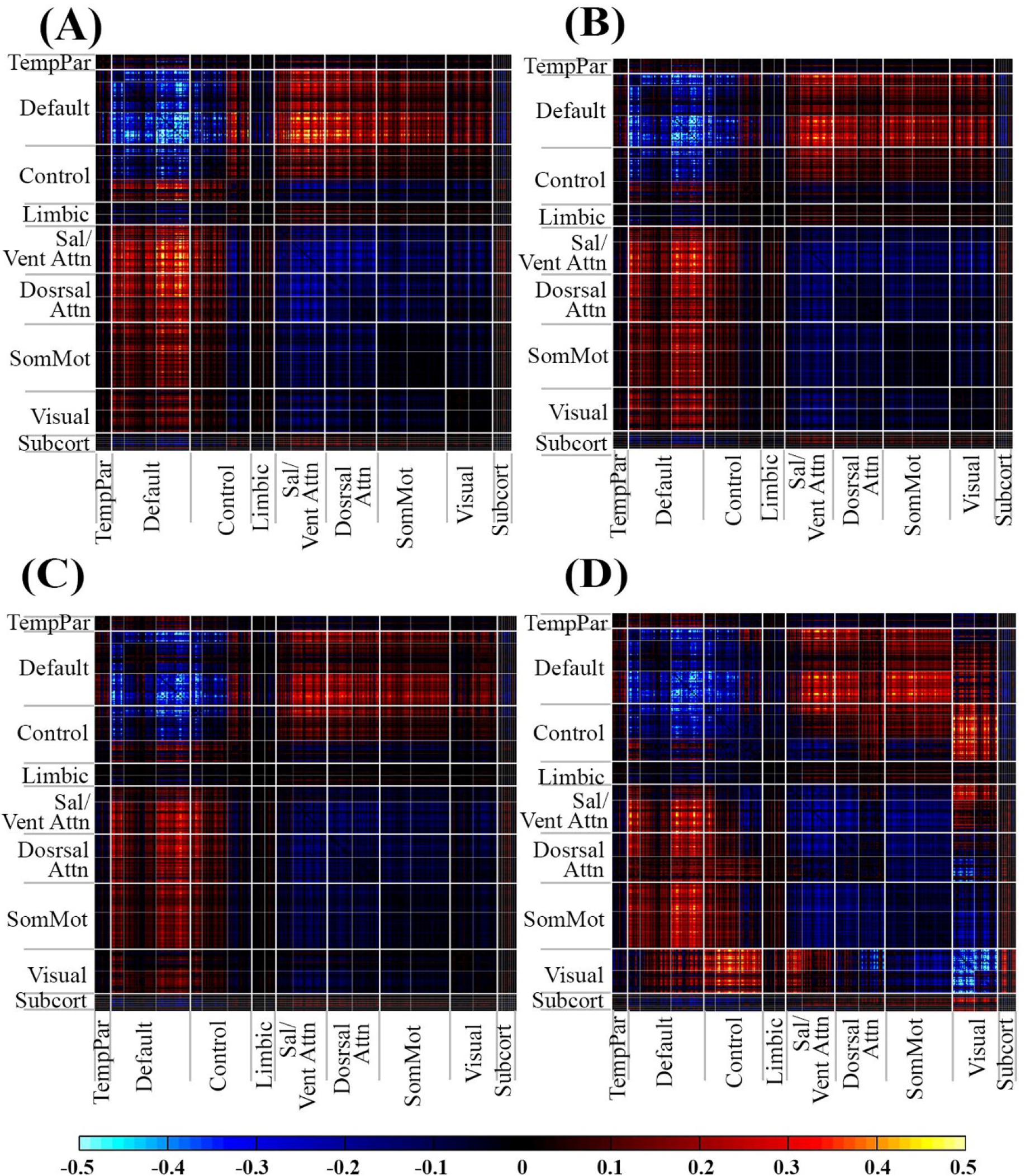
FC changes from removing the first common COBE component from the (A) second run, (B) third run and (C) fourth run of the HCP data. FC changes were mostly restricted to the default network and its connectivity with other networks. (D) FC changes from removing the first and second common COBE components from the first rs-fMRI run. FC changes mostly involved the default and visual networks, and their interactions with other networks.

### COBE improves behavioral prediction

The 419 × 419 FC matrices from NO-COBE, COBE-1000, COBE-2111 and COBE-3222 were utilized for cross-validated prediction 58 behavioral measures across cognition, personality and emotion (see Methods). The 20-fold cross-validation resulted in 20 prediction accuracies for each behavior measure.

Figure 5 shows the prediction accuracies averaged across all behavioral measures. COBE-2111 performed the best with an average prediction accuracy r = 0.179 ± 0.015 (mean ± std). Compared with NO-COBE (r = 0.160 ± 0.015), COBE-2111 achieves a relative improvement of 11.7% (p < 0.0001). From COBE-2111 to COBE-3222, the prediction accuracy dropped to r = 0.163 ± 0.015. From COBE-3222 to COBE-4333 (not shown), the prediction accuracy dropped even further to r = 0.143 ± 0.014, which confirmed our decision not to explore more components.

Table S1 reports the prediction accuracies for each behavioral measure averaged across 20 folds. Figure 6 shows the prediction accuracies of all 58 behavioral measures for NO-COBE and COBE-2111. COBE-2111 achieved an average relative improvement of 16.5% over NO-COBE (when relative improvement was computed for each behavioral measure and averaged across all behavioral measures).

We also repeated the analyses using partial correlations, instead of Pearson’s correlation. COBE-2111 again achieved the best prediction accuracies (Figure S2). Compared with NO-COBE, COBE-2111 achieved a relative improvement of 13.6% (p < 0.0001).

**Figure 5.**
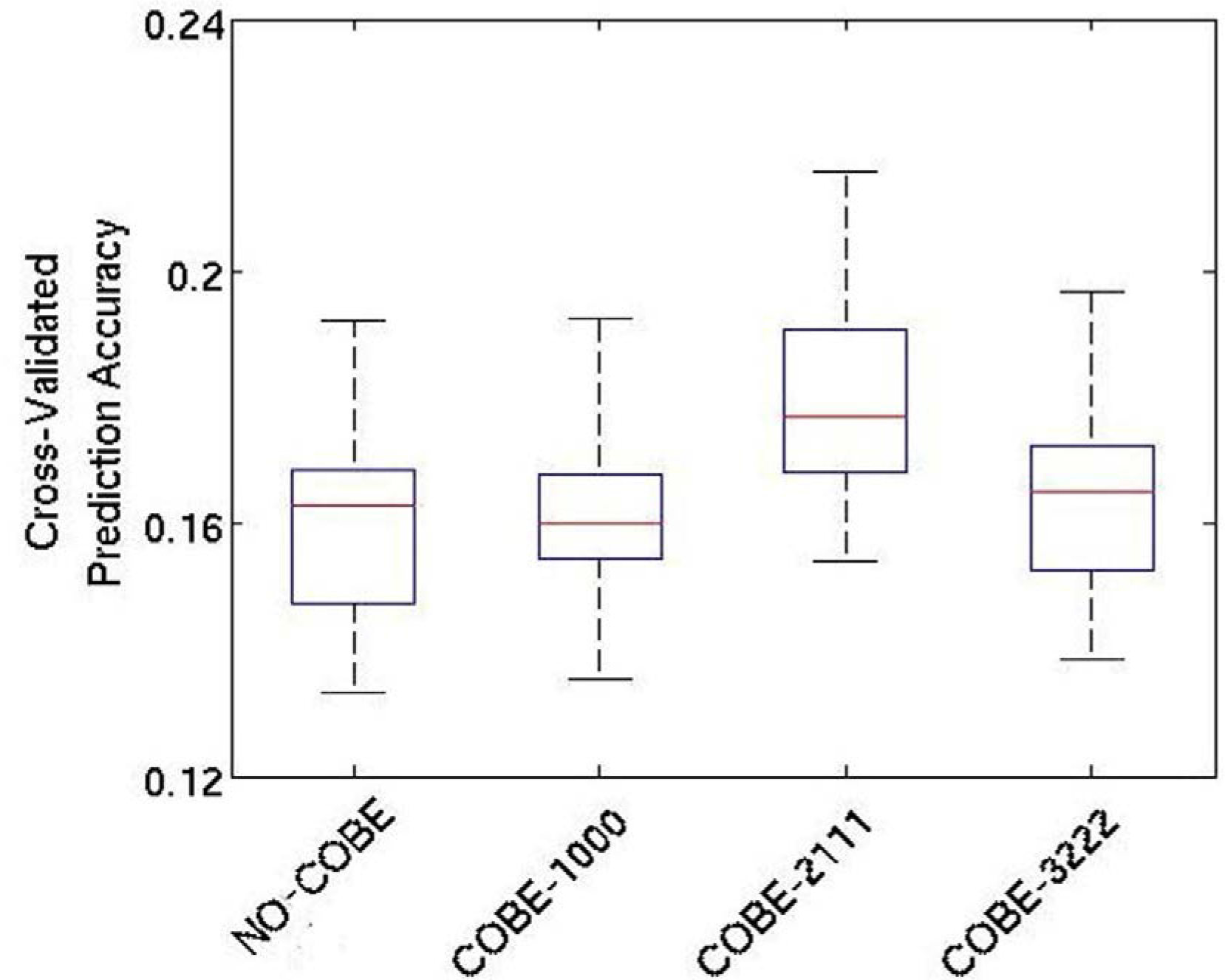
Cross-validated prediction accuracy (averaged across 58 behavioral measures) for NO-COBE, COBE-1000, COBE-2111 and COBE-3222. Functional connectivity was computed using Pearson’s correlation. COBE-2111 has the highest prediction accuracy.

**Figure 6.**
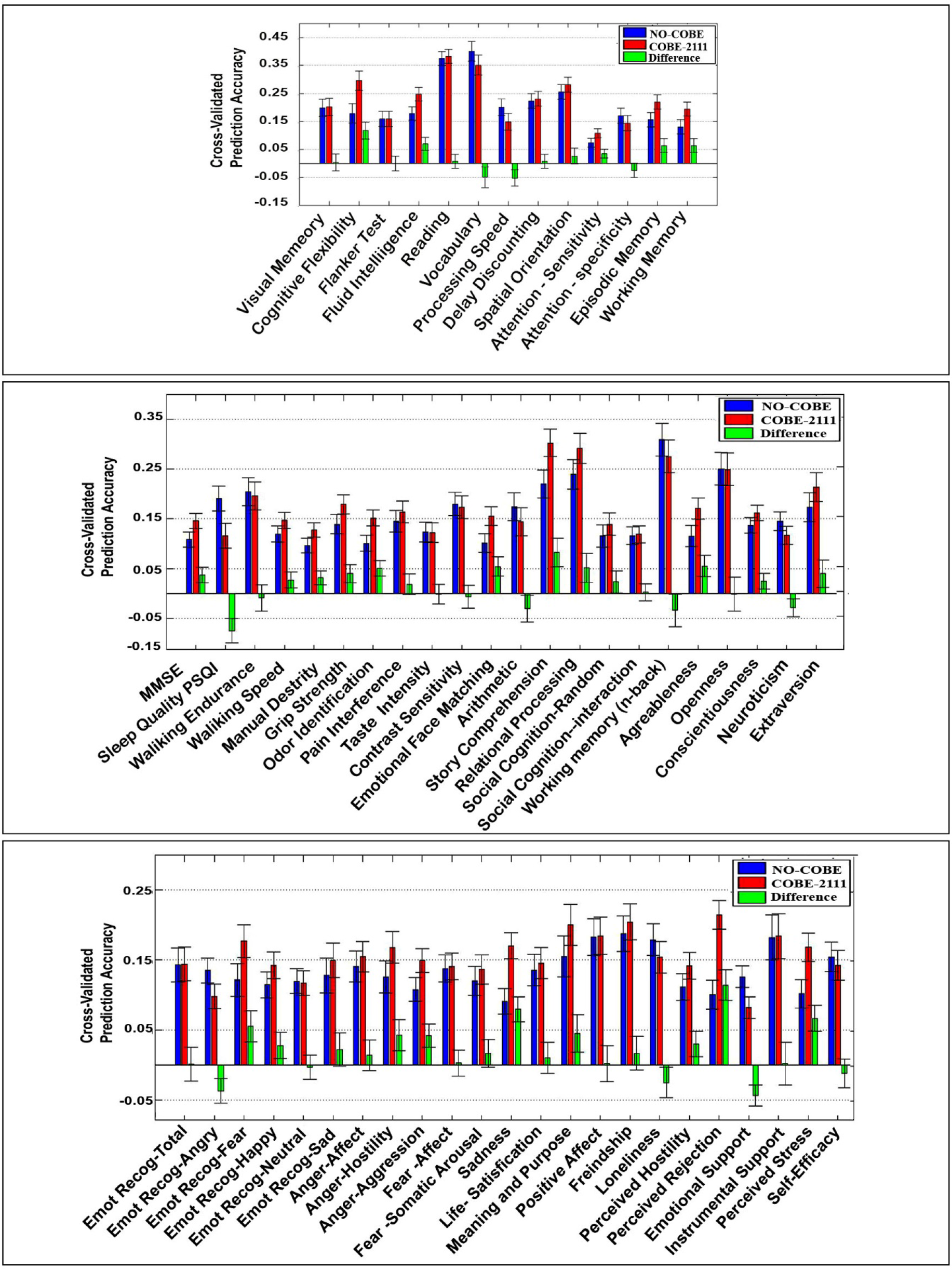
Cross-validated prediction accuracies (correlation) for NO-COBE and COBE-2111 for 58 behavioral measures across cognition, personality and emotion. Note the difference in y-axis scales across the three panels. COBE-2111 achieved an average relative improvement of 16.5% over NO-COBE

## Discussion

In this paper, we investigated whether the removal of common rs-fMRI components (that were shared across participants) could improve RSFC-based behavioral prediction. To this end, the COBE technique (Zhou et al., 2016a) was applied to decompose each HCP rs-fMRI run into a common subspace shared by all participants and individual-specific subspaces. We found that the first common COBE component from the first run was unique to that run. On the other hand, the second common COBE component from the first run was highly similar to the first common COBE component of the remaining three runs. By removing the first and second common COBE components from the first HCP run, and the first common COBE component from the remaining three HCP runs, behavioral prediction improved by 11.7% (averaged across 58 HCP behavioral measures).

Resting-state brain activity and functional connectivity are influenced by a large number of factors (Patriat et al., 2013; Yan et al., 2013; Rondinoni et al., 2011; Rondinoni et al., 2013; Tagliazuscchi et al., 2014; Gorgolewski et al., 2014; Laumann et al., 2015; Ong et al., 2015; Bijsterbosch et al., 2017; Power et al., 2017), including experimental conditions (e.g., fixation or eyes open rest, length of scan, etc), environment (e.g., scanner noise, temperature, etc), physiology (e.g., respiration and heart rate variability), brain state (e.g., caffeine intake, scanner anxiety, sleepiness) and content of self-generated thoughts (e.g., imagery, future related, etc).

In the case of the HCP data, Bijsterbosch and colleagues have noted run-specific (state-specific) differences, whereby early sensory-motor networks exhibited higher rs-fMRI amplitude in the second run of each of two scan days, with the first run of the first scan day exhibiting the lowest rs-fMRI amplitude. Our analyses also revealed run-specific effects: the first common COBE component of the first rs-fMRI run was unique and spatially localized in the peripherical portion of the visual cortex. Given that HCP participants were instructed to fixate on a bright cross-hair, one might speculate that this might yield strong effects in the visual cortex in the first run, which dissipated in subsequent runs as participants habituated to the “fixation task”.

Once the visual common component unique to the first HCP run was removed, the next common COBE component in the first run and the first common COBE component from the remaining three runs echoed the default network with particularly strong loadings on the posterior cingulate and precuneus, and also weak loadings on the inferior frontal gyrus, lateral temporal cortex, inferior parietal lobe and medial prefrontal cortex (Gusnard and Raichle, 2001; Raichle et al., 2001; Buckner et al., 2008). The default network is involved in self-generated thought (e.g., autobiographical memory, prospective thinking, etc), which is the dominant cognitive process during the resting-state (Spreng et al., 2009; Smallwood 2013; Andrews-Hanna et al., 2014). Furthermore, the posterior cingulate and precuneus are considered to be one of the core regions of the default network (Andrews-Hanna et al., 2010; Leech and Sharp, 2013) and plays a pivotal role in mediating the intrinsic activity throughout the default mode network (Fransson and Marrelec, 2008). Since the default network supports the generation of self-generated thoughts during the resting-state, it might not seem surprising that the default network is one of the common COBE component. However, inter-individual variation in the nature and content of self-generated thoughts can influence the resulting patterns of brain activity during rest (Gorgolewski et al., 2014; Wang et al., 2018). Therefore, it is somewhat surprising that the default network is one of the common COBE components that is shared across participants.

In the original COBE paper (Zhou et al., 2016a), COBE was shown to be useful in multiple datasets. For example, COBE was applied to 2856 face images from 68 individuals taken under different conditions (e.g., pose, illumination) to extract two common COBE components. By visual inspection, the common COBE components appeared to reflect illumination direction. Since illumination direction was not useful for face recognition, removing the common components improved recognition accuracies. Similarly, the common COBE components estimated in this study (Figure 2) might represent components not important for prediction, so their removal improved behavioral prediction. However, unlike face images (Zhou et al., 2016a), inferring the biological meaning of the components is tricky. While the common COBE components appeared biologically plausible (e.g., mapping to visual or default network regions), the actual biological mechanisms remain unclear and is a topic for future work.

## Conclusions

In this work, we decomposed participants’ rs-fMRI data into two components: a common (group-level) subspace and individual-specific subspaces. We found run-specific (state-specific) effects that were shared across participants. When common rs-fMRI signals were removed, the resulting RSFC yielded improved behavioral prediction in the Human Connectome Project.

## Acknowledgement

This work was supported by Singapore MOE Tier 2 (MOE2014-T2-2-016), NUS Strategic Research (DPRT/944/09/14), NUS SOM Aspiration Fund (R185000271720), Singapore NMRC (CBRG/0088/2015), NUS YIA and the Singapore National Research Foundation (NRF) Fellowship (Class of 2017). Our research also utilized resources provided by the Center for Functional Neuroimaging Technologies, P41EB015896 and instruments supported by 1S10RR023401, 1S10RR019307, and 1S10RR023043 from the Athinoula A. Martinos Center for Biomedical Imaging at the Massachusetts General Hospital. Our computational work was partially performed on resources of the National Supercomputing Centre, Singapore (https://www.nscc.sg). Data were provided by the Human Connectome Project, WU-Minn Consortium (Principal Investigators: David Van Essen and Kamil Ugurbil; 1U54MH091657) funded by the 16 NIH Institutes and Centers that support the NIH Blueprint for Neuroscience Research; and by the McDonnell Center for Systems Neuroscience at Washington University.

## References

Amico, Enrico, and Joaquín Goñi, 2018, “The quest for identifiability in human functional connectomes.” Sci Rep 8, 1, 8254.

Andrews-Hanna, Jessica R., Jonathan Smallwood, and R. Nathan Spreng, 2014, “The default network and self-generated thought: component processes, dynamic control, and clinical relevance.” Ann N Y Acad Sci, 1316, 1, 29–52.

Andrews-Hanna, J.R., Reidler, J.S., Sepulcre, J., Poulin, R. and Buckner, R.L., 2010. Functional-anatomic fractionation of the brain’s default network. Neuron, 65(4), 550–562.

Bijsterbosch, J., Harrison, S., Duff, E., Alfaro-Almagro, F., Woolrich, M., Smith, S., 2017. Investigations into within-and between-subject resting-state amplitude variations. Neuroimage, 159, 57–69.

Bertolero MA, Yeo BTT, Bassett DS, D’Espositor M. 2018, Strong hubs facilitate network modularity and cognitive performance. Nat. Hum. Behavi.

Biswal, B., Zerrin Yetkin, F., Haughton, V.M., Hyde, J.S., 1995. Functional connectivity in the motor cortex of resting human brain using echo-planar MRI. Magn Reson Med, 34, 537–541.

Bouckaert, R.R., Frank, E., 2004. Evaluating the replicability of significance tests for comparing learning algorithms. Pacific-Asia Conference on Knowledge Discovery and Data Mining. Springer pp. 3–12.

Buckner, R., Andrews-Hanna, J., Schacter, D., 2008, The brain’s default network: anatomy, function, and relevance to disease. Ann NY Acad Sci, 1124: 1–38.

Buckner, R.L., Krienen, F.M. and Yeo, B.T., 2013. Opportunities and limitations of intrinsic functional connectivity MRI. Nat Neurosci, 16 (7), 832.

Burgess, G.C., Kandala, S., Nolan, D., Laumann, T.O., Power, J.D., Adeyemo, B., Harms, M.P., Petersen, S.E., Barch, D.M., 2016. Evaluation of denoising strategies to address motion-correlated artefacts in resting-state functional magnetic resonance imaging data from the human connectome project. Brain Connect, 6, 669–680.

Cole, M.W., Bassett, D.S., Power, J.D., Braver, T.S., Petersen, S.E., 2014. Intrinsic and task-evoked network architectures of the human brain. Neuron, 83, 238–251.

Dubois, J., Galdi, P., Han, Y., Paul, L., & Adolphs, R. 2018. Resting-State Functional Brain Connectivity Best Predicts the Personality Dimension of Openness to Experience. Personal Neurosci, 1, E6.

Finn, E. S., Shen, X., Scheinost, D., Rosenberg, M. D., Huang, J., Chun, M. M., … & Constable, R. T. 2015. Functional connectome fingerprinting: identifying individuals using patterns of brain connectivity. Nat Neurosci, 18(11), 1664.

Finn, E.S., Constable, R.T., 2016. Individual variation in functional brain connectivity: implications for personalized approaches to psychiatric disease. Dialogues Clin Neurosci, 18, 277.

Fox, M.D., Raichle, M.E., 2007. Spontaneous fluctuations in brain activity observed with functional magnetic resonance imaging. Nat Rev Neurosci, 8, 700.

Fransson, P., Marrelec, G., 2008. The precuneus/posterior cingulate cortex plays a pivotal role in the default mode network: Evidence from a partial correlation network analysis. Neuroimage 42, 1178–1184.

Friedman, J., Hastie, T., Tibshirani, R., 2010. Regularization paths for generalized linear models via coordinate descent. J. Stat. Softw., 33, 1.

Fischl, B., Salat, D. H., Busa, E., Albert, M., Dieterich, M., Haselgrove, C., & Montillo, A. 2002. Whole brain segmentation: automated labeling of neuroanatomical structures in the human brain. Neuron, 33(3), 341–355.

Gratton, C., Laumann, T.O., Nielsen, A.N., Greene, D.J., Gordon, E.M., Gilmore, A.W., Nelson, S.M., Coalson, R.S., Snyder, A.Z., Schlaggar, B.L. and Dosenbach, N.U., 2018. Functional brain networks are dominated by stable group and individual factors, not cognitive or daily variation. Neuron, 98 (2), pp.439–452.

Greene, A.S., Gao, S., Scheinost, D. and Constable, R.T., 2018. Task-induced brain state manipulation improves prediction of individual traits. Nat. Commun., 9(1), 2807.

Gorgolewski, K.J., Lurie, D., Urchs, S., Kipping, J.A., Craddock, R.C., Milham, M.P., Margulies, D.S. and Smallwood, J., 2014. A correspondence between individual differences in the brain’s intrinsic functional architecture and the content and form of self-generated thoughts. PloS one, 9(5), 97176.

Glasser, M.F., Sotiropoulos, S.N., Wilson, J.A., Coalson, T.S., Fischl, B., Andersson, J.L., Xu, J., Jbabdi, S., Webster, M., Polimeni, J.R., 2013. The minimal preprocessing pipelines for the Human Connectome Project. Neuroimage 80, 105–124.

Griffanti, L., Salimi-Khorshidi, G., Beckmann, C.F., Auerbach, E.J., Douaud, G., Sexton, C.E., Zsoldos, E., Ebmeier, K.P., Filippini, N., Mackay, C.E., 2014. ICA-based artefact removal and accelerated fMRI acquisition for improved resting state network imaging. Neuroimage 95, 232–247.

Gusnard, D.A., Raichle, M.E., 2001. Searching for a baseline: functional imaging and the resting human brain. Nat Rev Neurosci, 2, 685.

Hampson, M., Driesen, N.R., Skudlarski, P., Gore, J.C., Constable, R.T., 2006. Brain connectivity related to working memory performance. J. Neurosci. 26, 13338–13343.

Jenkinson, M., Bannister, P., Brady, M., & Smith, S. 2002. Improved optimization for the robust and accurate linear registration and motion correction of brain images. Neuroimage, 17(2), 825–841.

Kong, R., Li, J., Orban, C., Sabuncu, M.R., Liu, H., Schaefer, A., Sun, N., Zuo, X.-N., Holmes, A., Eickhoff, S.B., 2018. Spatial Topography of Individual-Specific Cortical Networks Predicts Human Cognition, Personality and Emotion. Cereb Cortex, bhy123

Krienen, F.M., Yeo, B.T., Buckner, R.L., 2014. Reconfigurable task-dependent functional coupling modes cluster around a core functional architecture. Phil. Trans. R. Soc. B 369, 20130526.

Laumann, T.O., Gordon, E.M., Adeyemo, B., Snyder, A.Z., Joo, S.J., Chen, M.Y., Gilmore, A.W., McDermott, K.B., Nelson, S.M., Dosenbach, N.U. and Schlaggar, B.L., 2015. Functional system and areal organization of a highly sampled individual human brain. Neuron, 87(3), 657–670.

Leech, R., Sharp, D.J., 2013. The role of the posterior cingulate cortex in cognition and disease. Brain 137, 12–32.

Li, J., Kong, R., Liegeois, R., Orban, C., Sun, N., Holmes, A.J., Sabuncu, M.R., Ge, T., Yeo, B.T.T., 2018. Global Signal Regression Strengthens Association between Resting-State Functional Connectivity and Behavior. Under Review.

Mennes, M., Kelly, C., Zuo, X.-N., Di Martino, A., Biswal, B.B., Castellanos, F.X., Milham, M.P., 2010. Inter-individual differences in resting-state functional connectivity predict task-induced BOLD activity. Neuroimage 50, 1690–1701.

Mejia AF, Nebel MB, Shou H, Crainiceanu CM, Pekar JJ, Mostofsky S, Caffo B, Lindquist MA. 2015. Improving reliability of subject-level resting-state fMRI parcellation with shrinkage estimators. NeuroImage. 15;112:14–29.

Nadeau, C., Bengio, Y., 2000. Inference for the generalization error. Advances in neural information processing systems, pp. 307–313.

Ong, J.L., Kong, D., Chia, T.T., Tandi, J., Yeo, B.T. and Chee, M.W., 2015. Co-activated yet disconnected—Neural correlates of eye closures when trying to stay awake. Neuroimage, 118, 553–562.

Patriat, R., Molloy, E.K., Meier, T.B., Kirk, G.R., Nair, V.A., Meyerand, M.E., Prabhakaran, V., Birn, R.M., 2013. The effect of resting condition on resting-state fMRI reliability and consistency: a comparison between resting with eyes open, closed, and fixated. Neuroimage 78, 463–473.

Power, J.D., Plitt, M., Laumann, T.O. and Martin, A., 2017. Sources and implications of whole-brain fMRI signals in humans. Neuroimage, 146, 609–625.

Power, J. D., Barnes, K. A., Snyder, A. Z., Schlaggar, B. L., & Petersen, S. E. 2012. Spurious but systematic correlations in functional connectivity MRI networks arise from subject motion. Neuroimage, 59 (3), 2142–2154.

Raichle, M.E., MacLeod, A.M., Snyder, A.Z., Powers, W.J., Gusnard, D.A., Shulman, G.L., 2001. A default mode of brain function. Proc Natl Acad Sci U S A, 98, 676–682.

Rondinoni, C., dos Santos, A.C., Salmon, C.E.G., 2011. Effect of the scanner background noise on the resting brain networks detected by functional magnetic resonance imaging. Rev Bras Med Esporte, 5, 93–98.

Rondinoni, C., Amaro Jr, E., Cendes, F., Dos Santos, A.C. and Salmon, C.E.G., 2013. Effect of scanner acoustic background noise on strict resting-state fMRI. Braz J Med Biol Res, 46(4), 359–367.

Rosenberg, M. D., Finn, E. S., Scheinost, D., Papademetris, X., Shen, X., Constable, R. T., & Chun, M. M. 2016. A neuromarker of sustained attention from whole-brain functional connectivity. Nat Neurosci, 19(1), 165.

Spreng, R.N., Mar, R.A. and Kim, A.S., 2009. The common neural basis of autobiographical memory, prospection, navigation, theory of mind, and the default mode: a quantitative meta-analysis. J Cog Neurosci, 21(3), 489–510.

Smallwood, J., 2013. Distinguishing how from why the mind wanders: a process–occurrence framework for self-generated mental activity. Psychol bull, 139(3), 519.

Salimi-Khorshidi, G., Douaud, G., Beckmann, C.F., Glasser, M.F., Griffanti, L., Smith, S.M., 2014. Automatic denoising of functional MRI data: combining independent component analysis and hierarchical fusion of classifiers. Neuroimage 90, 449–468.

Schaefer, A., Kong, R., Gordon, E.M., Laumann, T.O., Zuo, X.-N., Holmes, A.J., Eickhoff, S.B., Yeo, B., 2017. Local-global parcellation of the human cerebral cortex from intrinsic functional connectivity MRI. Cereb. Cortex, 1–20.

Shirer, W.R., Ryali, S., Rykhlevskaia, E., Menon, V. and Greicius, M.D., 2012. Decoding subject-driven cognitive states with whole-brain connectivity patterns. Cereb. Cortex, 22 (1), 158–165.

Siegel, J.S., Mitra, A., Laumann, T.O., Seitzman, B.A., Raichle, M., Corbetta, M., Snyder, A.Z., 2016. Data quality influences observed links between functional connectivity and behavior. Cereb. Cortex 27, 4492–4502.

Smith, S.M., Beckmann, C.F., Andersson, J., Auerbach, E.J., Bijsterbosch, J., Douaud, G., Duff, E., Feinberg, D.A., Griffanti, L., Harms, M.P., 2013. Resting-state fMRI in the human connectome project. Neuroimage, 80, 144–168.

Smith, S.M., Fox, P.T., Miller, K.L., Glahn, D.C., Fox, P.M., Mackay, C.E., Filippini, N., Watkins, K.E., Toro, R., Laird, A.R., 2009. Correspondence of the brain’s functional architecture during activation and rest. Proc Natl Acad Sci U S A, 106, 13040–13045.

Smith, S.M., Nichols, T.E., Vidaurre, D., Winkler, A.M., Behrens, T.E., Glasser, M.F., Ugurbil, K., Barch, D.M., Van Essen, D.C., Miller, K.L., 2015. A positive-negative mode of population covariation links brain connectivity, demographics and behavior. Nat. Neurosci. 18, 1565.

Tagliazucchi, Enzo, and Helmut Laufs. 2014. Decoding wakefulness levels from typical fMRI resting-state data reveals reliable drifts between wakefulness and sleep. Neuron. 82, 695–708.

Tavor, I., Jones, O.P., Mars, R., Smith, S., Behrens, T., Jbabdi, S., 2016. Task-free MRI predicts individual differences in brain activity during task performance. Science, 352, 216–220.

Van Essen, D.C., Ugurbil, K., Auerbach, E., Barch, D., Behrens, T., Bucholz, R., Chang, A., Chen, L., Corbetta, M., Curtiss, S.W., 2012. The Human Connectome Project: a data acquisition perspective. Neuroimage, 62, 2222–2231.

Wang, C., Ong, J.L., Patanaik, A., Zhou, J. and Chee, M.W., 2016. Spontaneous eyelid closures link vigilance fluctuation with fMRI dynamic connectivity states. Proc Natl Acad Sci U S A, 113(34), 9653–9658.

Wang, H.T., Bzdok, D., Margulies, D., Craddock, C., Milham, M., Jefferies, E. and Smallwood, J., 2018. Patterns of thought: population variation in the associations between large-scale network organisation and self-reported experiences at rest. NeuroImage, 176, 518–527.

Yan, C.-G., Craddock, R.C., Zuo, X.-N., Zang, Y.-F., Milham, M.P., 2013. Standardizing the intrinsic brain: towards robust measurement of inter-individual variation in 1000 functional connectomes. Neuroimage 80, 246–262.

Yeo, B. T., Krienen, F.M., Sepulcre, J., Sabuncu, M.R., Lashkari, D., Hollinshead, M., Roffman, J.L., Smoller, J.W., Zöllei, L., Polimeni, J.R., 2011. The organization of the human cerebral cortex estimated by intrinsic functional connectivity. J Neurophysiol 106, 1125–1165.

Yeo, B.T., Krienen, F.M., Eickhoff, S.B., Yaakub, S.N., Fox, P.T., Buckner, R.L., Asplund, C.L., Chee, M.W., 2015a. Functional specialization and flexibility in human association cortex. Cereb Cortex 25, 3654–3672.

Yeo, B.T., Tandi, J., Chee, M.W., 2015b. Functional connectivity during rested wakefulness predicts vulnerability to sleep deprivation. Neuroimage 111, 147–158.

Zhou, G., Cichocki, A., Zhang, Y., Mandic, D.P., 2016a. Group component analysis for multiblock data: Common and individual feature extraction. IEEE Trans. Neural Netw. Learn. Syst, 2426–2439.

Zhou, G., Zhao, Q., Zhang, Y., Adalı, T., Xie, S., Cichocki, A., 2016b. Linked component analysis from matrices to high-order tensors: Applications to biomedical data. Proc. IEEE 104, 310–331.

Zou, H., Hastie, T., 2005. Regularization and variable selection via the elastic net. J R Stat Soc Series B Stat Methodol, 67, 301–320.

