## Supplemental for "Individual-Specific fMRI-Subspaces Improve Functional Connectivity Prediction of Behavior"

### Supplemental Results

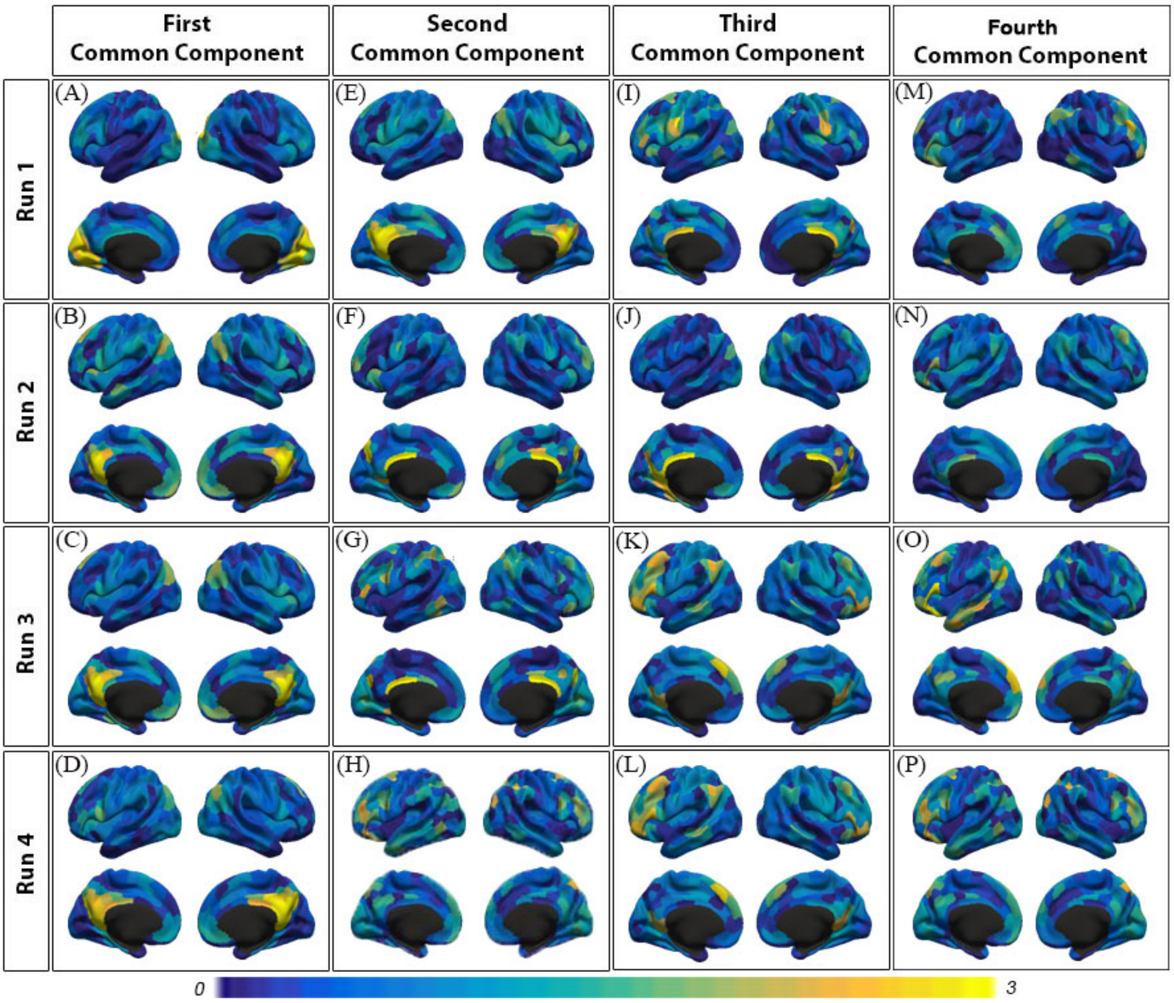


Figure S1. Spatial map of four common components shared across subjects (N = 803) in each of the four rs-fMRI runs. Observe that the first component of the first run (Figure S1A) was unique to only that run. Instead, the second component of the first run and the first component of the remaining runs were highly similar (r = 0.83). The third component of the first run and the second component of the remaining runs were also similar (r = 0.42). The fourth component of the first run and the third component of the remaining runs were not very similar (r = 0.06).

***Table S1***. Average cross-validated accuracies (Pearson's correlation) across 20 folds using NO-COBE, COBE-1000, COBE-2111 and COBE-3222 for 58 HCP behavioral measures. The highest accuracy for each behavioral measure is bolded.

| S. No | Behavior Category | Behavior Name | HCP Field Name | Average of 20-fold cross-validated accuracies | | | |
|---|---|---|---|---|---|---|---|
| | Cognition | | | NO-COBE | COBE-1000 | COBE-2111 | COBE-3222 |
| 1 | | Visual memory | PicSeq_Unadj | .19 | .16 | **.20** | .19 |

| | | | | | | | |
|---|---|---|---|---|---|---|---|
| 2 | | Cognitive flexibility | CardSort_Unadj | .17 | .24 | **.29** | .24 |
| 3 | | Flanker test | Flanker_Unadj | .15 | .15 | .15 | .15 |
| 4 | | Fluid intelligence | PMAT24_A_CR | .17 | .21 | **.24** | .18 |
| 5 | | Reading | ReadEng_Unadj | .37 | .38 | **.38** | .36 |
| 6 | | Vocabulary | PicVocab_Unadj | **.4** | .34 | .35 | .34 |
| 7 | | Processing speed | ProcSpeed_Unadj | **.2** | .14 | .14 | .14 |
| 8 | | Delay discounting | DDisc_AUC_40K | .22 | .21 | **.23** | .17 |
| 9 | | Spatial orientation | VSPLOT_TC | .25 | .27 | **.28** | .26 |
| 10 | | Attention – Sensitivity | SCPT_SEN | .07 | .10 | .10 | **.13** |
| 11 | | Attention – Specificity | SCPT_SPEC | **.16** | .15 | .14 | **.16** |
| 12 | | Episodic Memory | IWRD_TOT | .15 | **.21** | **.21** | .18 |
| 13 | | Working Memory | ListSort_Unadj | .13 | .14 | **.19** | **.19** |
| 14 | Personality | Cognitive Status | MMSE_Score | .1 | .08 | **.14** | .09 |
| 15 | | Sleep quality (PSQI) | PSQI_Score | **.19** | .15 | .11 | .16 |
| 16 | | Walking endurance | Endurance_Unadj | .20 | .18 | .19 | **.22** |
| 17 | | Walking Speed | GaitSpeed_Unadj | .11 | .12 | **.14** | .12 |
| 18 | | Manual Dexterity | Dexterity_Unadj | .09 | .11 | **.12** | .11 |
| 19 | | Grip Strength | Strength_Unadj | .13 | .11 | **.17** | .11 |
| 20 | | Odor identification | Odor_Unadj | 0.1 | .1 | **.15** | .11 |
| 21 | | Pain Interference Survey | PainInterf_Tscore | .14 | .16 | **.16** | .13 |
| 22 | | Taste intensity | Taste_Unadj | .12 | **.15** | .12 | .11 |
| 23 | | Contrast Sensitivity | Mars_Final | **0.17** | .13 | **.17** | .13 |
| 24 | | Emotional face matching | Emotion_Task_Face_Acc | .10 | .12 | **.15** | .11 |

| | | | | | | | |
|---|---|---|---|---|---|---|---|
| 25 | | Arithmetic | Language_Task_Math_Avg_Difficulty_Level | **.17** | **.17** | .14 | .16 |
| 26 | | Story comprehension | Language_Task_Story_Avg_Difficulty_Level | .21 | .28 | **.30** | .26 |
| 27 | | Relational processing | Relational_Task_Acc | .23 | .25 | **.29** | .27 |
| 28 | | Social Cognition – random | Social_Task_Perc_Random | .11 | .12 | **.13** | .11 |
| 29 | | Social Cognition – interaction | Social_Task_Perc_TOM | .11 | .10 | .11 | **.14** |
| 30 | | Working Memory (n-back) | WM_Task_Acc | **.30** | .26 | .27 | .23 |
| 31 | | Agreeableness | NEOFAC_A | .11 | .15 | **0.17** | 0.15 |
| 32 | | Openness | NEOFAC_O | **.25** | .24 | .24 | .23 |
| 33 | | Conscientiousness | NEOFAC_C | .13 | .12 | **.16** | .12 |
| 34 | | Neuroticism | NEOFAC_N | **.14** | .08 | .11 | .09 |
| 35 | | Extraversion | NEOFAC_E | .17 | .16 | **.21** | .18 |
| 36 | Emotion | Emotion Recognition – Total | ER40_CR | .14 | **.15** | .14 | .12 |
| 37 | | Emotion Recognition – Angry | ER40_ANG | **.13** | .09 | .09 | .10 |
| 38 | | Emotion Recognition – Fear | ER40_FEAR | .12 | .11 | **.17** | .11 |
| 39 | | Emotion Recognition – Happy | ER40_HAP | .11 | .11 | **.14** | .10 |
| 40 | | Emotion Recognition – Neutral | ER40_NOE | .11 | .10 | .11 | **.12** |
| 41 | | Emotion | ER40_SAD | .12 | .12 | **.14** | **.14** |

| | | | | | | | |
|---|---|---|---|---|---|---|---|
| | | Recognition – Sad | | | | | |
| 42 | | Anger – Affect | AngAffect_Unadj | .14 | .13 | **.15** | **.15** |
| 43 | | Anger – Hostility | AngHostil_Unadj | .12 | .13 | **.16** | .11 |
| 44 | | Anger – Aggression | AngAggr_Unadj | .10 | .14 | **.14** | .12 |
| 45 | | Fear – Affect | FearAffect_Unadj | .13 | .13 | **.14** | .12 |
| 46 | | Fear – Somatic Arousal | FearSomat_Unadj | .12 | **.15** | .13 | .13 |
| 47 | | Sadness | Sadness_Unadj | .09 | .14 | **.17** | .15 |
| 48 | | Life Satisfaction | LifeSatisf_Unadj | .13 | .13 | .14 | **.16** |
| 49 | | Meaning & Purpose | MeanPurp_Unadj | .15 | .16 | **.19** | .16 |
| 50 | | Positive Affect | PosAffect_Unadj | **.18** | .16 | **.18** | **.18** |
| 51 | | Friendship | Friendship_Unadj | .18 | .19 | **.21** | .19 |
| 52 | | Loneliness | Loneliness_Unadj | **.17** | .15 | .15 | .12 |
| 52 | | Perceived Hostility | PercHostil_Unadj | .11 | .13 | .14 | **.15** |
| 54 | | Perceived Rejection | PercReject_Unadj | .10 | .12 | **.21** | .14 |
| 55 | | Emotional Support | EmotSupp_Unadj | **.12** | .08 | .08 | .10 |
| 56 | | Instrument Support | InstruSupp_Unadj | **.18** | .15 | **.18** | .17 |
| 57 | | Perceived Stress | PercStress_Unadj | .10 | .11 | **.16** | .15 |
| 58 | | Self - Efficacy | SelfEff_Unadj | .15 | .13 | .14 | **.18** |

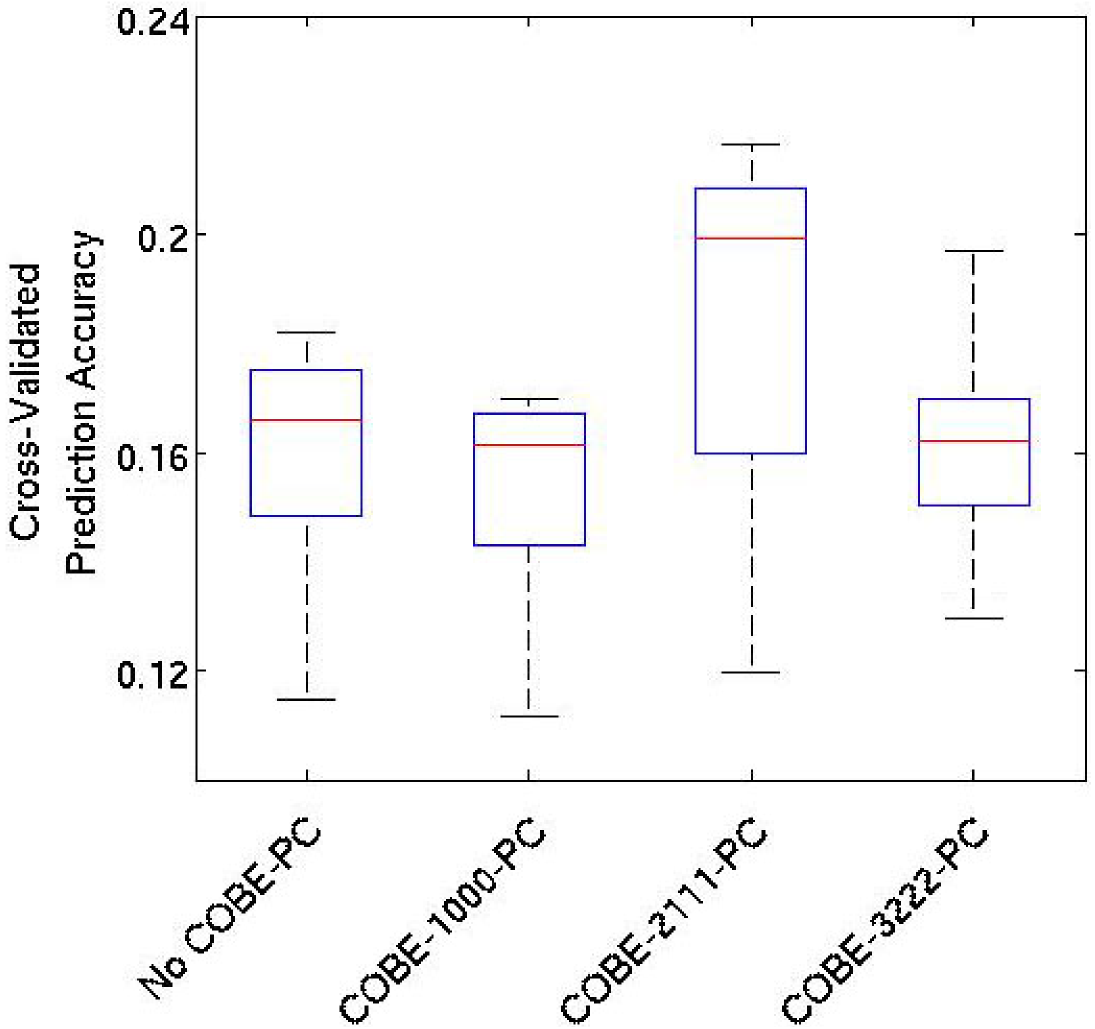


Figure S2. Cross-validated prediction accuracy (averaged across 58 behavioral measures) for NO-COBE-PC, COBE-1000-PC, COBE-2111-PC and COBE-3222-PC. Functional connectivity was computed using partial correlations. COBE-2111-PC has the highest prediction accuracy.
